# Endothelial iron homeostasis regulates BBB integrity via the HIF2α – Ve-cadherin pathway

**DOI:** 10.1101/2020.10.28.358473

**Authors:** Daniel Rand, Orly Ravid, Dana Atrakchi, Hila Israelov, Yael Bresler, Chen Shemesh, Liora Omesi, Sigal Liraz-Zaltsman, Fabien Gosselet, Taber S. Maskrey, Michal Schnaider Beeri, Peter Wipf, Itzik Cooper

## Abstract

The blood-brain barrier (BBB) serves as the guardian of the CNS, tightly regulating the movement of ions, molecules, and cells between the circulatory system and brain. This barrier is critical in maintaining brain homeostasis, allowing proper neuronal function and protecting the brain from injury and disease. Chronic and acute exposure to various chemicals lead to BBB breakdown through pathways that are also affected in neurological diseases. Therefore, we have created an in-vitro BBB injury model to gain a better understanding of the mechanisms controlling BBB integrity. This model exposes a co-culture of human stem-cell derived brain-like endothelial cells (BLEC) and brain pericytes that mimic the BBB, to the organophosphate paraoxon. This exposure results in rapid lipid peroxidation, initiating a ferroptosis-like process and leading to endothelium cell toxicity. Mitochondrial ROS formation (MRF) and increase in mitochondrial membrane permeability (MMP), which occur 8 - 10 h post paraoxon-induced injury, also trigger apoptotic cell death. Yet, these processes do not directly result in damage to barrier functionality since blocking them does not reverse the increased permeability. Looking for a crucial pathway affecting barrier functionality we analyzed the iron homeostasis in our model since the iron chelator, Desferal© (DFO) rescued endothelial cell viability. Upon BBB insult, the liable iron pool (LIP) is rapidly increased, preventing the increased expression of the stress related hypoxia-induced factor 2α (HIF2α) transcription factor. This results in a decrease in surface expression of the adherens junction and permeability master regulator protein, Ve-cadherin, ultimately damaging BBB integrity. Unlike the apoptosis inhibitor ZVAD that rescues BLEC from cell toxicity, yet exacerbates damage to the barrier functionality, DFO significantly decreases MRF and apoptosis subsequent to PX exposure, while also rescuing barrier integrity by inhibiting the liable iron pool increase, inducing HIF2α expression and preventing the degradation of Ve-cadherin on the cell surface. Moreover, the novel nitroxide JP4-039 significantly rescues both injury-induced endothelium cell toxicity and barrier functionality. Collectively, we have elucidated the cellular processes initiated by chemical injury to the endothelium barrier that result in cell toxicity; yet, inhibiting these processes does not necessarily protect BBB integrity which is regulated by the iron mediated HIF2α – Ve-Cadherin axis. DFO protects BBB integrity by inhibiting the injury-induced deregulation of this axis. Additionally, we have discovered a novel compound, JP4-039, that inhibits both damage to endothelium functionality and cell toxicity. Elucidating a regulatory pathway that maintains BBB integrity and discovering both a novel and an FDA approved compound that interfere with this pathway elucidates a potential therapeutic approach to protect the BBB degradation that is evident in many neurological diseases.

## Introduction

The blood–brain barrier (BBB) is composed of the capillaries of the central nervous system (CNS) which tightly regulate the movement of molecules, ions, and cells between the blood and the CNS ^1,2^. The BBB further protects the CNS by preventing neurotoxic plasma components, blood cells, and pathogens from entering the brain^3^. It is formed by a monolayer of tightly-sealed endothelial cells that make up the walls of the capillaries, and together with the closely associated pericytes and astrocytic end-feet processes control the restricted flow of compounds in and out of the brain, both through their paracellular junctions and by a limited array of transcellular vesicular and specialized transporters routes^4–6^.

BBB breakdown and dysfunction leads to leakage of harmful blood components into the CNS, cellular infiltration, and aberrant transport and clearance of molecules^3,6,7^. Alzheimer’s disease (AD), amyotrophic lateral sclerosis (ALS) and Parkinson disease (PD) are all associated with defective BBB function^8–12^. A hallmark of these neurodegenerative disorders is an excessive amount of reactive oxygen species (ROS), which is viewed as one of the potential common etiologies in these diseases^13–15^.

Imbalance between ROS production and antioxidant defenses results in excessive accumulation of ROS and leads to oxidative stress^16,17^. Oxidative stress results in cell membrane damage from lipid peroxidation, changes in protein structure and function due to protein degradation, and structural damage to DNA^18^. The brain is especially susceptible to oxidative stress due to its high oxygen demand, constituting 20% of body oxygen consumption ^19^. Additionally, redox-active metals such as iron exist abundantly in the brain and are actively involved in the formation and propagation of ROS. Lastly, high levels of polyunsaturated fatty acids are present in brain cell membranes and form substrates for lipid peroxidation^20^.

In the aging brain, a region-specific increase of total iron content is observed, probably triggered by inflammation, increased BBB permeability, redistribution of iron within the brain, and changes in iron homeostasis^21–23^. Changes in regional iron distribution have been demonstrated consistently in Neurodegenerative diseases^24,25^. Additionally, many Neurodegenerative diseases, such as Alzheimer’s and Parkinson diseases, are primarily characterized by a deposition of insoluble protein aggregates of which iron is an essential component^26–29^.

All these components seem to be interconnected, yet how they influence each other is still unclear. High iron levels, BBB permeability and oxidative stress are all biomarkers for neurodegenerative diseases^30–32^ yet it is still uncertain whether these are isolated events or an interlinked physiological cascade.

To understand the relationship and kinetics of these processes and their role in BBB breakdown in neurodegenerative diseases we established a BBB-injury model based on the organophosphate paraoxon (PX) and a stem-cell derived in-vitro human BBB system^33–37^ since there is a high correlation between neurodegenerative diseases and exposure to various chemicals and specifically to organophosphate pesticides^38–40^. Additionally, this exposure induces common effects that are predominant in neurodegenerative diseases, such as; direct damage to BBB integrity^41,42^ and ROS production among varied cell types, including endothelial cells^43–45^. Our model exhibited many of these common BBB cellular abnormalities including increased permeability, ROS formation, inflammation and cell death^36,37^.

In this work, we have shown that iron rather than oxidative stress, is responsible for the barrier functionality and elucidated the novel brain endothelial HIF2α – Ve-Cadherin pathway responsible for this damage. Furthermore, we have discovered potential therapeutic compounds– the iron chelator, Desferal© (DFO, an FDA-approved drug for the treatment of iron overload) and a novel nitroxide, JP4-039– that mitigate both damage to endothelium viability and functionality. This discovery can aid in clarifying the role of iron in BBB breakdown in neurodegenerative diseases, ultimately resolving the age-old paradox of whether iron is a cause or consequence of these diseases^46–48^ and provide a new therapeutic target for multiple brain diseases in which the BBB is impaired.

## Results

### Lipid peroxidation is rapidly induced in a BBB injury model

We have previously reported that neither antioxidants nor apoptosis inhibitors were able to rescue barrier functions in our BBB injury model^36,37^, suggesting the source of this damage is not a direct result of either process. Therefore, our goal was to elucidate the changes in molecular and signaling pathways responsible for the functional damage in the brain-like endothelial cells (BLEC) of the BBB injury model. Since high levels of oxidative stress were evident in this model, yet antioxidants such as TEMPOL were unable to rescue barrier integrity^36^, we aimed to ascertain whether inhibiting a specific type of oxidative stress would provide this outcome. Since lipid compositions and their physiochemical properties greatly influence barrier functions^49^ and intercellular lipids are a common target for oxidation by ROS^50,51^, we measured lipid peroxidation at the initial insult event. A rapid and significant increase in lipid peroxidation was observed (Fig. 1a). Additionally, we measured lipid peroxidation over a 24-hour period using a live-cell analysis system and found that lipid peroxides accumulated over time (Fig. 1b). The lipid peroxidation inhibitor Ferrostatin-1 (Fer-1) ^52^ abrogated this damage throughout the entire experiment (Fig. 1a and b). Cell death was abrogated in the presence of Fer-1, indicating that lipid peroxidation contributed to the increased cell toxicity (Fig. 1c). These results suggested that insult to BLEC might lead to ferroptosis for which Fer-1 is a classic and specific blocker^52^. The ferroptotic cell death entails two essential components: lipid peroxidation and accumulation of free iron ions^52,53^. We therefore treated cells with Desferal (DFO), an FDA-approved drug which acts as an iron chelator. Indeed, DFO significantly rescued cell toxicity (Fig. 1c). To further confirm the ferroptosis process, the absence of apoptosis needed to be established ^52,54^. The kinetics of apoptosis induction were measured with live cell imaging of caspase-3 activation for the duration of 24 hours subsequent to endothelium exposure to PX. We found that PX induced apoptosis in BLEC, and that the addition of Fer-1 did not abrogate this effect (Fig. 1d). Therefore, even though most of the classic ferroptosis signatures existed, it was inaccurate to classify this process as ferroptosis per se since ultimately, apoptosis occurred (Fig. 1d). We then asked whether this ferroptosis-like process could account for the impaired functionality observed in the BBB model; Even though Fer-1 abrogated cell death (Fig. 1c), it did not rescue barrier function from increased permeability (Fig. 1e). Collectively, these results suggested that there were at least two processes that occurred as a result of PX damage to the brain endothelial cells. Initially, the cells rapidly responded to the PX challenge by lipid peroxidation; however, after 12 hours, additional cell death pathways came into play, such as apoptosis. These seemed to be independent pathways, since BLEC underwent apoptosis even in the presence of Fer-1 (Fig. 1d). Additionally, neither ferroptosis nor apoptosis seemed to be solely accountable for increased BBB permeability since two classic blockers of these programmed cell death processes (Fer-1 and the pan caspase inhibitor ZVAD) significantly attenuated cell death but failed to rescue the damage to barrier permeability (Fig. 1e and ^36^).

**Fig. 1.**
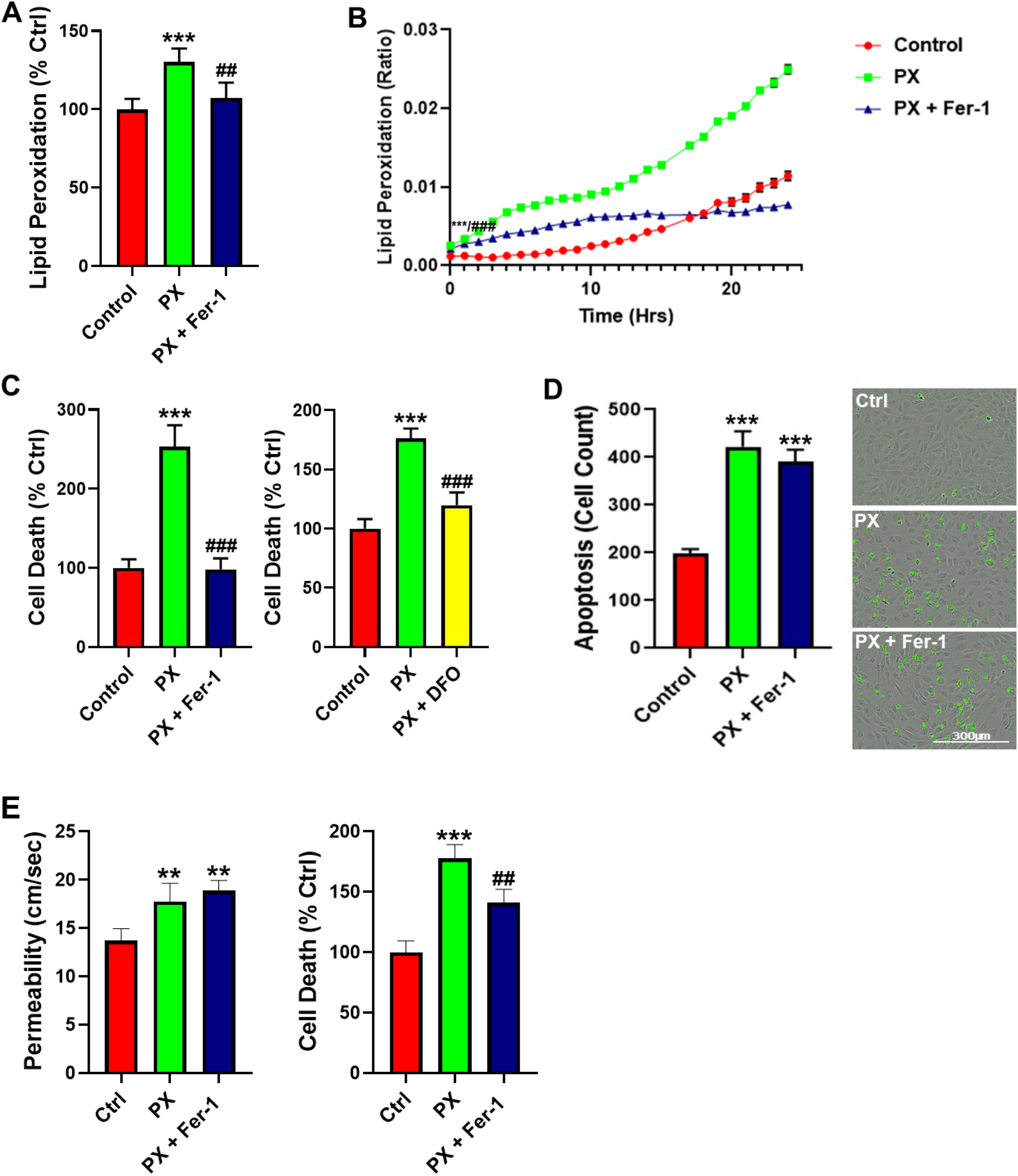
Inhibition of lipid peroxidation rescues cell viability but not barrier function. **A.** Cellular lipid peroxidation subsequent to PX treatment on a BLEC monolayer (PX: Paraoxon, 400 μM; Fer-1: Ferrostatin-1, 50 μM). Mean ± s.d. of n = 9-12 for each condition from 3 independent experiments, ***p ≤ 0.001 vs ctrl and ### p ≤ 0.01 vs PX. **B.** Cellular lipid peroxidation assessed by live cell imaging assay over a 24-hour period on a BLEC monolayer. Mean ± s.d. of n = 4-6 for each condition from a single experiment, ***p ≤ 0.001 vs ctrl and ### p ≤ 0.001 vs PX marks the first time point with a significant change. **C.** Cell death assessed by the release of lactate dehydrogenase (LDH) 24-hour post PX treatment on a BLEC monolayer. (PX: Paraoxon, 400 μM; Fer-1: Ferrostatin-1, 50 μM; DFO: Desferal 50μg/ml). Mean ± s.d. of n = 10-18 for each condition from 3 independent experiments, ***p ≤ 0.001 vs ctrl and ## p ≤ 0.001 vs PX. **D.** Apoptosis was measured with a caspase-3 activated fluorescent marker by live cell imaging 24-hour post PX treatment on a BLEC monolayer. Mean ± s.d. of n = 9-12 for each condition from 3 independent experiments, ***p ≤ 0.001 vs ctrl. **E.** Permeability of sodium fluorescein (NaF) was measured across the BBB in vitro model (from luminal to abluminal side) 24-hour post PX treatment (PX: Paraoxon, 900 μM; Fer-1: Ferrostatin-1, 50 μM). Cell death was assessed by sampling medium from the luminal side with the LDH assay. Mean ± s.d. of n = 9-15 for each condition from 3 independent experiments, **p ≤ 0.01 and ***p ≤ 0.001 vs ctrl and ## p ≤ 0.01 vs PX.

### Kinetics of mitochondrial ROS formation

Since Fer-1 failed to inhibit PX-induced apoptosis (Fig. 1d), we assumed that apoptosis was not a direct result of the initial lipid peroxidation. Since inhibition of several oxidative stress associated events in our model, such as lipid peroxidation (Fig. 1) and total cell ROS formation 36, did not rescue the BBB integrity, we next measured mitochondrial ROS production. Mitochondrial superoxide formation was significantly increased 9 hours after PX treatment in a time dependent manner (Fig. 2a). Since mitochondrial stress initiates the chain of events leading to apoptosis^55^, we treated the cells with ZVAD, which significantly decreased PX-induced mitochondrial ROS formation (MRF), while Fer-1 exacerbated it (Fig. 2a), suggesting that this organelle-specific ROS formation might be responsible for the chain of events leading to apoptosis, although not as a direct consequence of the initial lipid peroxidation. It has been reported that mitochondrial superoxide formation leads to a decrease in mitochondrial membrane potential and causes mitochondrial membrane permeation (MMP) ^56^ which is the next step in the chain of events terminating in apoptosis. We therefore performed a live cell imaging assay to measure the permeation of the mitochondrial membrane in cells treated with PX. At 15 hours post treatment, the permeation started to increase significantly in comparison to the control cells, increasing until 24 hours. ZVAD abrogated this effect and Fer-1 had no significant effect at 24 hours (Fig. 2b). Additionally, ZVAD prevented the activation of caspase-3 and reduced the basal apoptosis process of untreated cells after 24 hours (Fig. 2c). Concordantly, ZVAD nullified cell cytotoxicity throughout the 24-hour treatment (Fig. 2d). Together, these results suggest two chain of events and their kinetics in this model of injured endothelium; one results in apoptosis while the other leads to a ferroptosis-like process (Fig. 2e). However, even though these experiments clarified both chains of events and provided insights into their time frames, the source of functional damage in the brain endothelium had yet to be clarified.

**Fig. 2.**
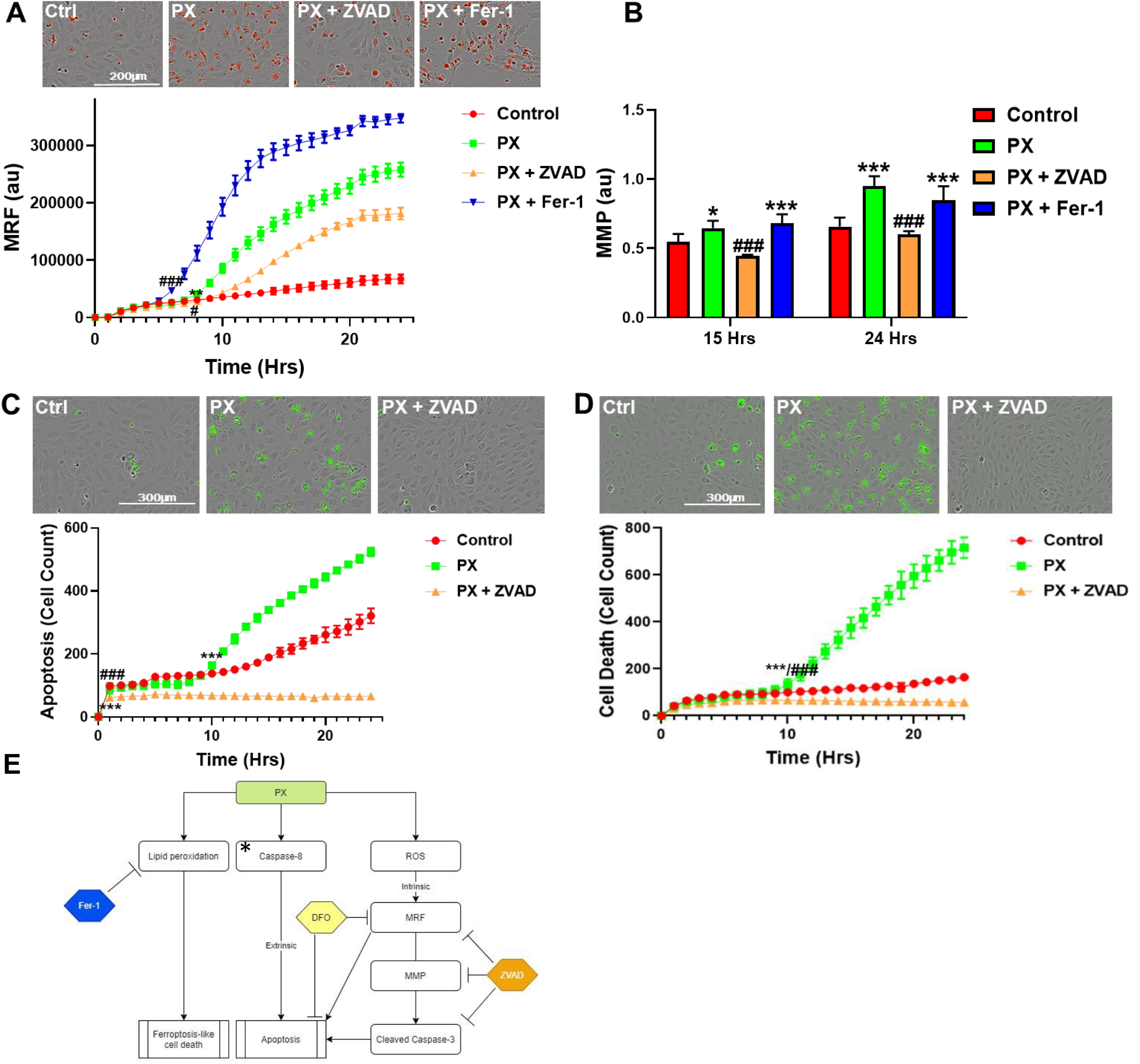
Apoptosis and mitochondrial stress are not a direct result of the early lipid peroxidation initiated by PX. **A.** Monolayers of BLEC were treated with PX and a combination of PX and ZVAD or Ferrostatin-1 for 24 hours (PX at 400 μM, ZVAD at 50 μM, Fer-1 at 50 μM). Live cell imaging of mitochondrial ROS formation was measured. Mean ± s.d. of n = 12-18 for each condition from 3 independent experiments, **p ≤ 0. 01, # p ≤ 0.05 and ### p ≤ 0.001 marks the first significant difference versus control and PX-treated cells respectively. Images captured 24 hours post treatment. **B.** Mitochondrial membrane permeability was examined by measuring its membrane depolarization with JC-1 staining over a 24-hour period on a BLEC monolayer. Mean ± s.d. of n = 12-18 for each condition from 3 independent experiments, *p≤ 0.05, ***p ≤ 0.001 and ### p ≤ 0.001 marks a significant difference versus control and PX-treated cells respectively. **C.** Live cell imaging of a caspase-3 activated fluorescent marker was used to measured apoptosis on a BLEC monolayer. Mean ± s.d. of n = 9-12 for each condition from 3 independent experiments, ***p ≤ 0.001 and ## p ≤ 0.001 marks the first time point with a significant change versus control and PX-treated cells respectively. Images captured 24 hours post treatment. **D.** Live cell imaging of cell death was examined by cytotoxicity fluorescent staining over a 24-hour period on a BLEC monolayer. Mean ± s.d. of n = 12-18 for each condition from 3 independent experiments, ***p ≤ 0.001 and ## p ≤ 0.001 marks the first time point with a significant change versus control and PX-treated cells respectively. Images captured 24 hours post treatment. **E.** Flowchart elucidating the molecular pathways initiated in our model.* We have recently reported that inhibiting caspase-8 decreased apoptosis in our model^37^. MRF, mitochondrial ROS formation; MMP, mitochondrial membrane permeability; au, arbitrary units

### The iron chelator DFO rescues both cell viability and functionality

Iron is essential for redox catalysis and bioenergetics; nevertheless, if not appropriately complexed, iron also plays a key role in the formation of toxic oxygen radicals that can attack many biomolecules^57^. Since several ROS processes were evident in our model and DFO rescued cells from PX-induced cell death (Fig. 1b), we wanted to further elucidate its mechanism. DFO rescued cytotoxicity throughout the 24-hour window (Fig. 3a). Since DFO is an iron chelator, we hypothesized that damage to BLEC deregulated the cellular iron homeostasis. Indeed, upon PX treatment, there was a significant increase in the liable iron pool within the cell, while DFO moderately but significantly rescued this effect (Fig. 3b). Unlike Fer-1, DFO did not rescue the cells exclusively through the novel ferroptosis-like process since it also significantly decreased apoptosis (Fig. 3c). Since apoptosis was inhibited, we next measured the effect of DFO on MMP, a hallmark of the apoptosis pathway^58^. Conversely, DFO exacerbated the damage to the mitochondrial membrane (Fig. 3d), suggesting that DFO did not affect apoptosis through the classic chain of events that are associated with the breakdown of the mitochondrial membrane and the release of pro-apoptotic proteins such as cytochrome C as discussed above (Fig. 2e). Therefore, we next measured the cellular ROS formation since the Fenton reaction is a catalytic process in which iron is oxidized by hydrogen peroxide, a product of mitochondrial oxidative respiration, ultimately leading to the creation of a highly toxic hydroxyl free radical which results in oxidative stress^59^. Surprisingly, DFO had no significant effect on cellular ROS formation (Fig. 3e). It has been reported that the Fenton reaction initiates the production of hydroxyl radicals within the mitochondria^60^, leading to oxidative stress. Indeed, DFO significantly decreased mitochondrial ROS formation (Fig. 3f), suggesting that DFO rescues cell viability through inhibition of mitochondrial superoxide formation. Conversely, DFO significantly increased the permeation of the mitochondrial membrane rather than preventing it (Fig. 3d) indicating that DFO does not rescue cell viability via the classic apoptosis pathway. Our next step was to test whether the ability of DFO to rescue cell viability in BLEC also extended to their functionality. DFO significantly rescued barrier function as manifested by preventing the increase in permeability and the decrease in transendothelial electrical resistance (TEER) in the BBB model exposed to PX (Fig. 3g). This confirms that DFO rescues both cell viability and barrier functionality. Collectively, these results highlight the therapeutic potential of addressing iron dyshomeostasis in the treatment of BBB damage since this approach has the dual effect of restoring loss of function as well as inhibiting cell toxicity in the endothelial cells of the BBB.

**Fig. 3.**
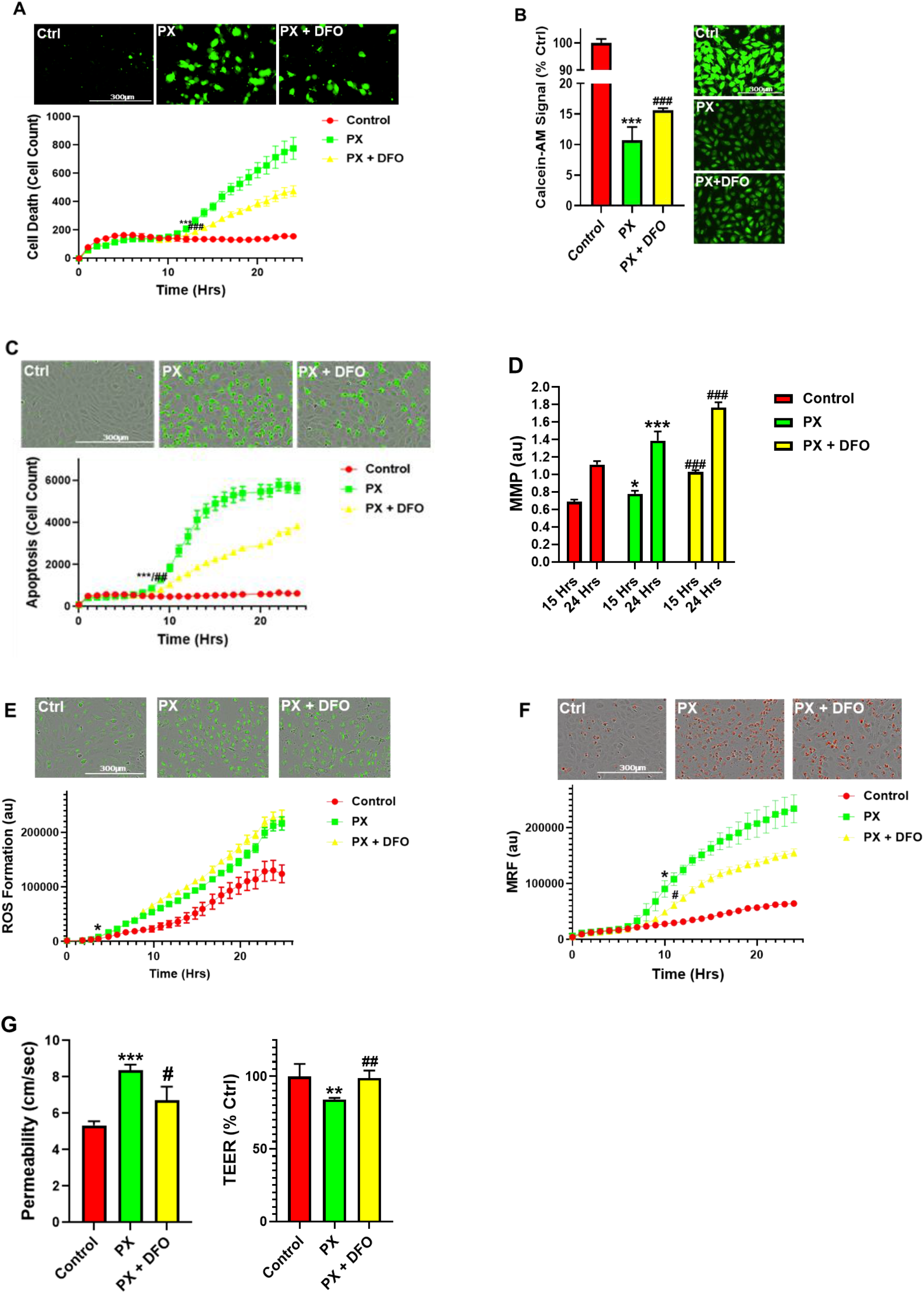
DFO diminishes cell toxicity and rescues barrier functions. **A.** Live cell imaging of cell toxicity measured over a period of 24 hours on a BLEC monolayer (PX: Paraoxon, 400 μM; DFO: Desferal, 50 μg/ml) mean ± s.d. of n = 20-30 for each condition from 5 independent experiments, ***p ≤ 0.001 and ### p ≤ 0.01 marks the first significant difference versus control and PX-treated cells respectively. Images captured 24 hours post treatment. **B.** Liable Iron Pool (LIP) was assessed by Calcein-AM on a BLEC monolayer. Its fluorescent signal negatively correlates with LIP. Mean ± s.d. of n = 10-12 for each condition from 2 independent experiments, ***p ≤ 0.001 and ## p ≤ 0.001 versus control and PX-treated cells respectively. Images captured 1-hour post treatment. **C.** Apoptosis was examined by live imaging of activated caspase-3 staining on a BLEC monolayer. Mean ± s.d. of n = 12-18 for each condition from 3 independent experiments, ***p ≤ 0.001 and ## p ≤ 0.01 marks the first significant difference versus control and PX-treated cells respectively. Images captured 24 hours post treatment. **D.** Mitochondrial membrane permeability was evaluated with the use of live imaging by measuring the mitochondrial membrane depolarization with JC-1 staining on a BLEC monolayer. Mean ± s.d. of n = 12-18 for each condition from 3 independent experiments, *p ≤ 0.05, **p ≤ 0.01 and ### p ≤ 0.001 marks the first significant difference versus control and PX-treated cells respectively. Images captured 24 hours post treatment. **E.** Live cell imaging of cellular ROS formation was measured with CellRox staining on a BLEC monolayer. Mean ± s.d. of n = 12-18 for each condition from 3 independent experiments, *p ≤ 0.05 marks the first significant difference versus control. Images captured 24 hours post treatment. **F.** Live cell imaging of mitochondrial ROS formation was measured with the Mitosox marker on a BLEC monolayer. Mean ± s.d. of n = 16-24 for each condition from 4 independent experiments, *p ≤ 0. 05 and # p ≤ 0.05 marks the first significant difference versus control and PX-treated cells respectively. Images captured 24 hours post treatment. **G.** Permeability of NaF was measured across the BBB in-vitro model (from luminal to abluminal side) 24 hours post PX treatment. Using the same model, TEER (Ω.cm2.) was also measured. Mean ± s.d. of n = 12-15 for each condition from 3 independent experiments, ***p ≤ 0.001, **p ≤ 0.01, # p ≤ 0.05 and ## p ≤ 0.01 versus control and PX-treated cells respectively. MRF, mitochondrial ROS formation; MMP, mitochondrial membrane permeability; au, arbitrary units

### DFO rescues BBB integrity by protecting Ve-cadherin through regulation of HIF2α

We next wanted to elucidate the molecular mechanism of DFO by which it rescues BBB integrity. Since no correlation was found between lipid peroxidation, ROS formation or apoptosis and barrier integrity, we decided to investigate the effect of DFO on adherens and tight junctions which are responsible for the paracellular tightness of the BBB. Out of all the adherens and tight junctional proteins we evaluated; Ve-cadherin expression was the most affected by exposure to PX in-vivo and in-vitro^36,37^. Additionally, Ve-cadherin is known to be a master regulator of permeability in BBB endothelial cells^61^. We therefore measured Ve-cadherin expression in cells exposed to a combined treatment of PX and DFO. Using western blot and immunocytochemistry, DFO was shown to abrogate the reduction in total Ve-cadherin protein expression (Fig. 4a). To investigate a connection between the increased liable iron pool and Ve-cadherin down regulation, we measured the levels of the iron regulated transcription factor; hypoxia induced factor 2 alpha (HIF2α). This transcription factor has been implicated as a Ve-cadherin regulator via different pathways including directly regulating Ve-cadherin expression or expression of other proteins that stabilize it such as vascular endothelial protein tyrosine phosphatase (Ve-PTP) ^62,63^. While there was no significant effect on HIF2α expression in the injury model, addition of DFO significantly increased HIF2α expression levels in comparison to Control and PX-treated cells (Fig 4b). These results translated exclusively to the functional level since the HIF2α inhibitor abrogated the rescue effect of DFO in both permeability and TEER but not cell viability (Fig 4c). This strengthened the hypothesis that the liable iron pool is responsible for the damage to BBB integrity since this pattern replicated itself when PX was exchanged with ferrous ammonium sulfate (FAS) to directly increase the liable iron pool^64^ while not inducing cell toxicity (Fig 4d). HIF2α is a transcription factor that specifically induces Ve-cadherin expression independently of hypoxia^63^. We therefore measured mRNA levels in BLEC exposed to PX. Surprisingly, even though HIF2α inhibitor had an adverse yet not significant effect on Ve-cadherin levels, DFO did not rescue mRNA decrease in response to insult (Fig 4e), suggesting that DFO rescued Ve-cadherin expression post transcriptionally. We then measured the effect of HIF2α on Ve-cadherin whole cell protein levels and found that even though DFO prevented the decrease in Ve-cadherin protein expression, HIF2α inhibitor did not abrogate this effect (Fig 4f). Conversely, Ve-cadherin cell surface expression reflected a clear pattern of DFO rescue and HIF2α inhibitor attenuation of this rescue effect (Fig 4g). To ascertain that this functional pathway is independent of the ROS associated apoptotic pathways mentioned earlier (Fig. 2e), we measured MRF. As expected, HIF2α inhibitor had no significant effect on DFO rescue (Fig 4h), strengthening the hypothesis that endothelium functionality is not regulated by ROS related processes in our model. Collectively, these results suggest that HIF2α regulates BBB integrity through maintaining Ve-cadherin on the cell surface. Damage to the BBB increases the cellular liable iron pool, resulting in a decrease in HIF2α-mediated Ve-cadherin cell surface expression, while DFO restores this expression, resulting in the functional rescue of the BBB endothelial cells.

**Fig. 4.**
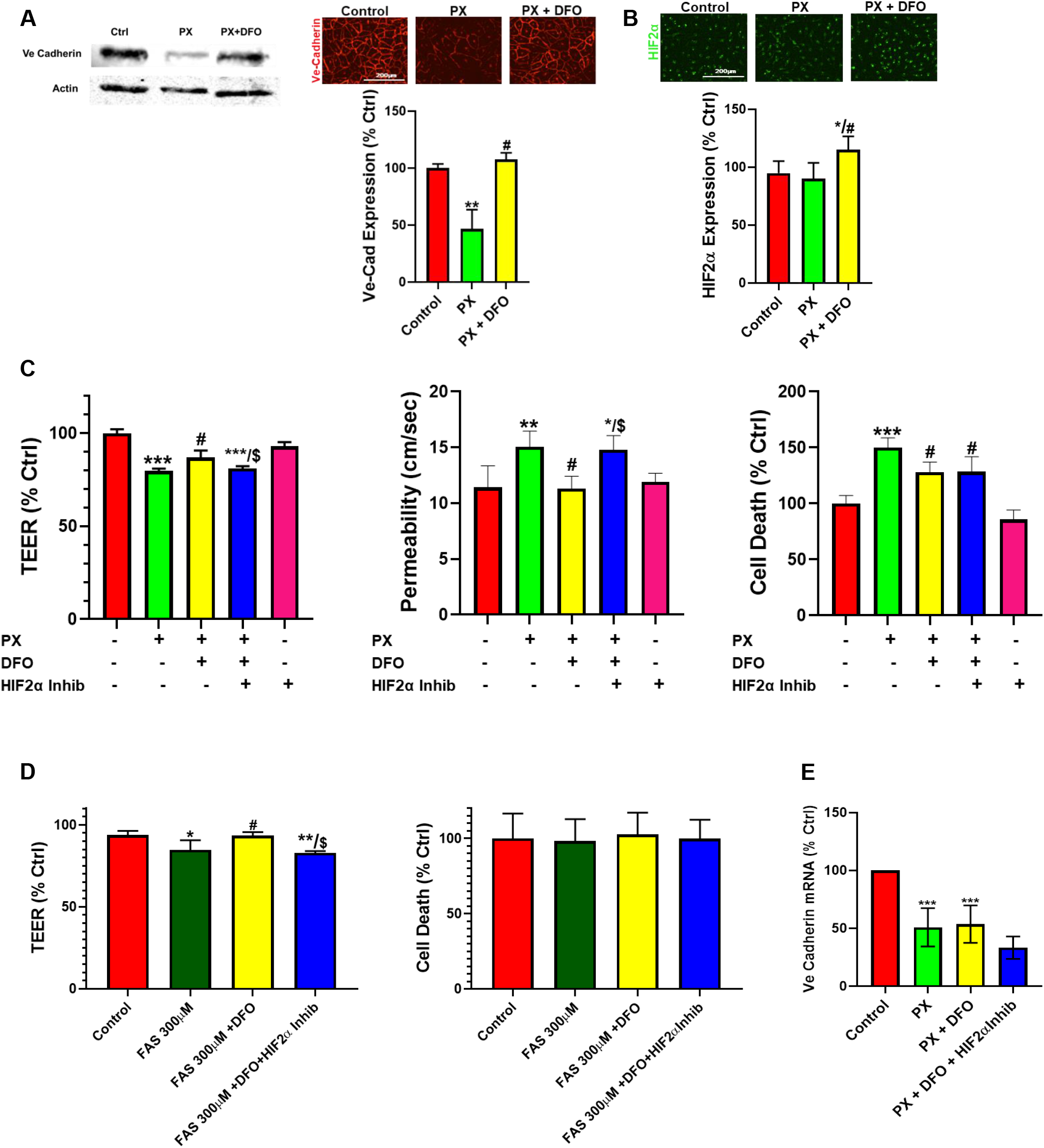
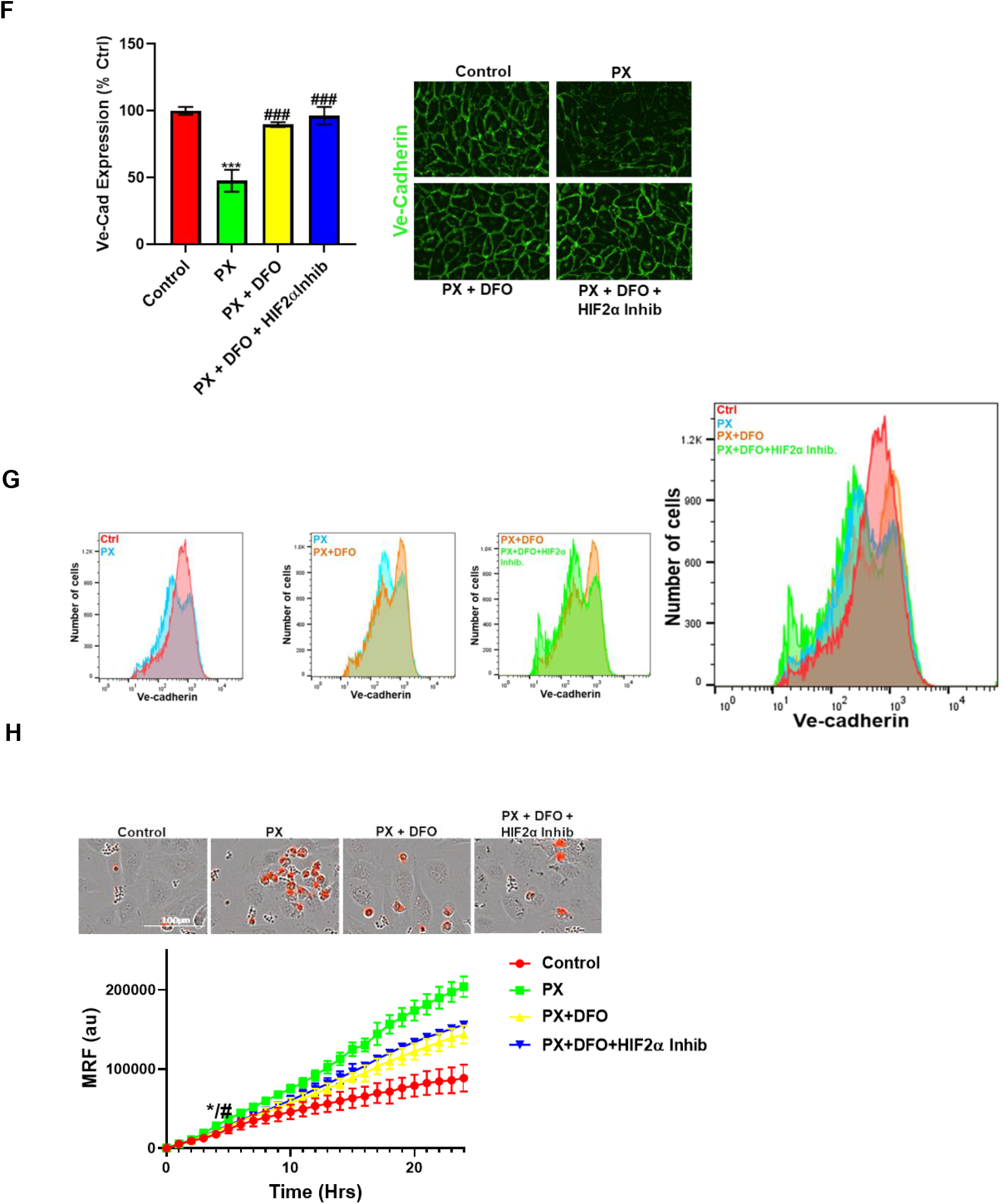
DFO rescues BBB Permeability via a HIF2α-Ve-cadherin pathway. **A.** Ve-Cadherin expression was measured in a BLEC monolayer with western blot and immunocytochemistry intensity quantification in a BLEC monolayer treated with PX +/− DFO (PX: Paraoxon 400 μM, DFO: Desferal 50 μg/ml) for 24 hour, n = 16-24 for each condition from 4 independent experiments, **p ≤ 0.01 and # p ≤ 0.05 versus control and PX-treated cells respectively. **B.** Immunofluorescence staining and quantification of HIF2α in a BLEC monolayer treated with PX (400 μM) ± DFO (50 μg/ml) for 24 hour, n = 16-24 for each condition from 4 independent experiments, *p ≤ 0.05 and # p ≤ 0.05 versus control and PX-treated cells respectively. **C.** Permeability of NaF was measured across the BBB in-vitro model (from luminal to abluminal side) 24 hours post PX (900 μM) ± DFO (50 μg/ml) ± HIF2α inhibitor (10 μM) treatment. Using the same model, TEER (Ω.cm^2^.) was also measured. Cell death was assessed by sampling medium from the luminal side with the LDH assay. Mean ± s.d. of n = 9-15 for each condition from 3 independent experiments, ***p ≤ 0.001, ** p ≤ 0.01 and ** p ≤ 0.05 vs control, # p ≤ 0.05 and vs PX and $ p ≤ 0.05 vs PX + DFO. **D.** TEER (Ω.cm^2^.) was measured across the BBB in-vitro model 12 hours post FAS (300 μM) ± DFO (50 μg/ml) ± HIF2α inhibitor (10 μM) treatment. Cell death was assessed by sampling medium from the luminal side with the LDH assay. Mean ± s.d. of n = 6-10 for each condition from 2 independent experiments, **p ≤ 0.01 and * p ≤ 0.05 vs control, # p ≤ 0.05 vs PX and $ p ≤ 0.05 vs PX + DFO. **E.** RT-PCR quantification of Ve-Cadherin mRNA in a BLEC monolayer treated with PX (400 μM) ± DFO (50 μg/ml) ± HIF2α inhibitor (10 μM) for 24 hours. Mean ± s.d. of n=5-6 for each condition from 6 biological repeats and 3 technical repeats per each n, ***p ≤ 0.001 versus control. **F.** Immunofluorescence staining and quantification of Ve-Cadherin in a BLEC monolayer treated with PX (400 μM) ± DFO (50 μg/ml) ± HIF2α inhibitor (10 μM) for 24 hours. Mean ± s.d. of n = 16-24 for each condition from 4 independent experiments, ***p ≤ 0.001 and ### p ≤ 0.001 versus control and PX-treated cells respectively. **G.** FACS histograms of live single cells from a BLEC monolayer treated with PX (400 μM) ± DFO (50 μg/ml) ± HIF2α inhibitor (10 μM) for 24 hours. Cells were stained for Ve-Cadherin. Immunostaining was performed without fixation and permeabilization (surface staining), n = 3 for each condition from 3 independent experiments. Median decrease of 32% with PX treatment (p < 0.001), DFO rescue of 22% (p < 0.01) and HIF2α inhibitor attenuated DFO rescue by 34% (p < 0.01). **H.** Live cell imaging of mitochondrial ROS formation was measured with Mitosox staining on a BLEC monolayer. Mean ± s.d. of n = 16 for each condition from 3 independent experiments, *p≤0.05 and # p≤ 0.05 marks the first significant difference versus control and PX-treated cells respectively. Images captured 24 hours post treatment.

### A new chemical entity, JP4-039, rescues barrier functions

Since the cellular iron homeostasis is a precisely and delicately regulated process that is present throughout all cells in the body^65,66^, we aimed at identifying another agent that does not elicit such a profound effect on the cellular iron levels. Like DFO, this compound should have the potential to mitigate the synergistic effects of increased cell death and decreased cell functionality induced in the damaged BBB. We found that the novel nitroxide, JP4-039 (Fig. 5a), significantly rescued permeability in our injury model (Fig. 5b) and inhibited TEER decrease (Fig. 5c). JP4-039 also inhibited cell death as recorded using the LDH assay (Fig. 5d). The cytotoxicity and apoptosis kinetics were measured in the presence of JP4-039. Both processes were initially inhibited at 8 to 10 hours and up to 24 hours post treatment (Fig. 5e and 5f). We have previously shown that JP4-039 is an antioxidant and electron scavenger that targets the mitochondria. Moreover, JP4-039 is also an inhibitor of erastin and RSL-3 induced ferroptotic cell death^67^. Indeed, JP4-039 significantly inhibited MRF increase (Fig. 5g) in PX treated cells. Collectively, these results suggested that the novel nitroxide, JP4-039, offers a therapeutic potential to treat BBB damage, since this compound has the dual effect of rescuing both the viability and integrity of the endothelial cells.

**Fig. 5.**
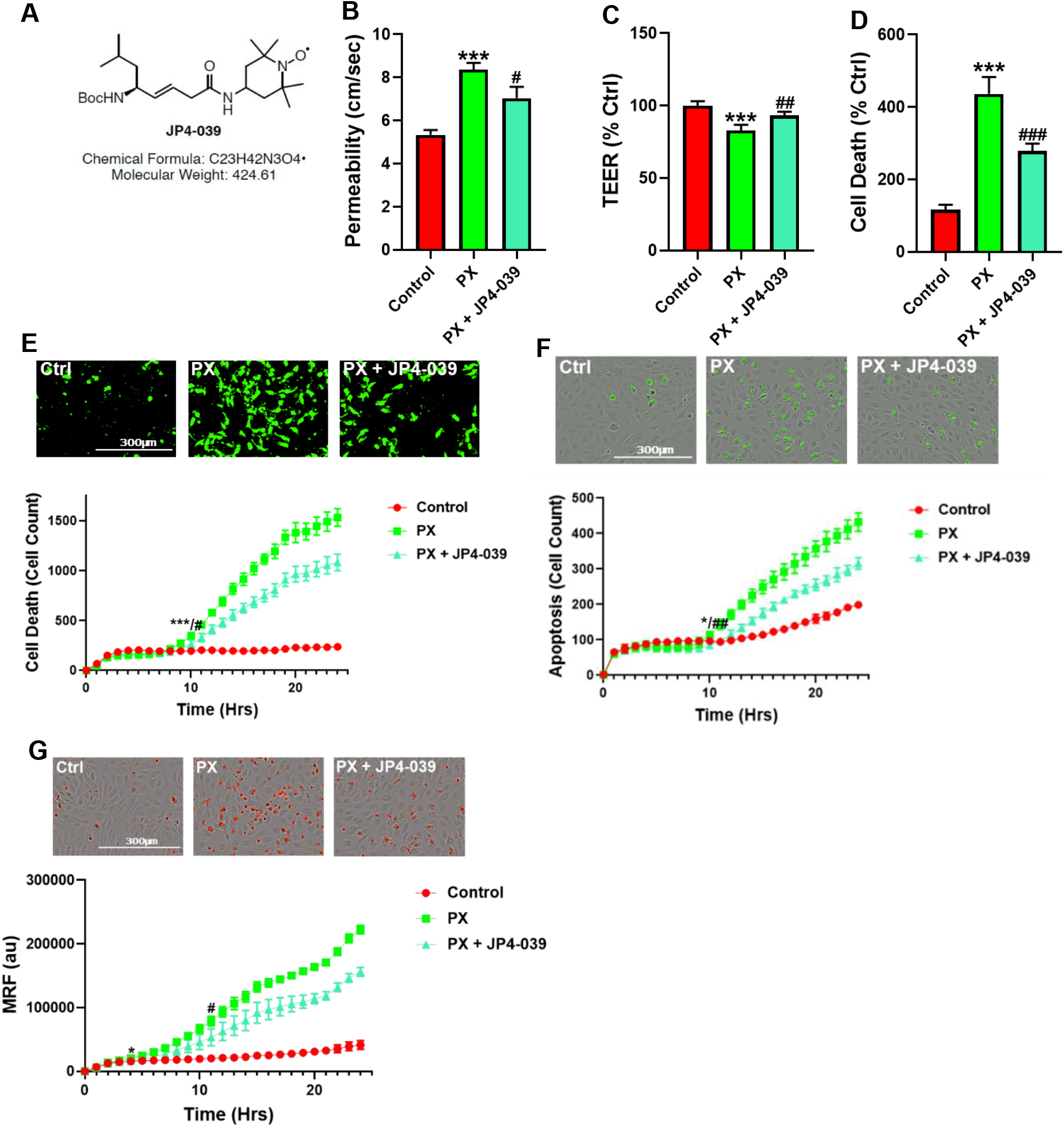
JP4-039 rescues PX-induced toxicity and permeability increase in the BBB injury model. **A.** A chemical formula diagram of JP4-039. **B.** Permeability of NaF was measured across the BBB in-vitro model (from luminal to abluminal side) 24 hours post PX (900 μM) ± JP4-039 (25 μM) treatment. Mean ± s.d. of n = 12-20 for each condition from 4 independent experiments, ***p≤0.001 and #p≤ 0.05 versus control and PX-treated cells respectively. **C.** Using the same model, TEER was also measured. Mean ± s.d. of n = 12-20 for each condition from 4 independent experiments, ***p ≤ 0.001 and ## p ≤ 0.01 versus control and PX-treated cells respectively. **D.** Cell death in a BLEC monolayer was assessed by measuring LDH from the cell culture media (PX at 400 μM, JP4-039 at 624 nM). Mean ± s.d. of n = 9-18 for each condition from 3 independent experiments, ***p ≤ 0.001 and ## p ≤ 0.001 versus control and PX-treated cells respectively. **E.** Live cell imaging of cell toxicity was measured over a period of 24 hours on a BLEC monolayer. Mean ± s.d. of n = 16-24 for each condition from 4 independent experiments, ***p ≤ 0.001 and # p ≤ 0.05 marks the first significant difference versus control and PX-treated cells respectively. Images captured 24 hours post treatment. **F.** Live cell imaging of apoptosis was examined by activated caspase 3 staining on a BLEC monolayer. Mean ± s.d. of n = 16-24 for each condition from 4 independent experiments, *p ≤ 0.05 and ### p ≤ 0.01 marks the first significant difference versus control and PX-treated cells respectively. Images captured 24 hours post treatment. **G.** Live cell imaging of Mitochondrial ROS formation with Mitosox staining on a BLEC monolayer. Mean ± s.d. of n = 20-30 for each condition from 5 independent experiments, *p ≤ 0. 05 and # p ≤ 0.05 marks the first significant difference versus control and PX-treated cells respectively. Images captured 24 hours post treatment.

## Discussion

In this study, we have used a human stem cell-derived BBB injury model to elucidate molecular mechanisms and their kinetics that are induced upon insult with an organophosphate toxin. We have shown that rapid lipid peroxidation is initiated, resulting in a ferroptosis-like process, which ultimately affects endothelium cell viability. Inhibition of this process with the classic lipid peroxidation and ferroptosis blocker Fer-1 abrogated endothelium cell death but had no effect on their reduced functionality. Additional processes came into effect around 10 hours post PX exposure, resulting in apoptosis. In our system apoptosis seems to be the result of the convergence of various pathways, rather than a classic single cell death pathway. The live cell imaging assays elucidated the kinetics of several classic molecular cascades resulting in apoptosis; we observed that 10 hours post insult there was a significant increase in mitochondrial superoxide formation which accumulated till 24 hours. This damage initiated mitochondrial membrane permeation, leading to caspase-3 activation and apoptosis. All these events leading to apoptosis were inhibited with the pan caspase inhibitor ZVAD. There is evidence that other pathways are initiated which result in apoptosis since the iron chelator DFO inhibited both mitochondrial superoxide formation and apoptosis yet exacerbated the increased mitochondrial membrane permeation. However, blocking all these ROS related stresses such as lipid peroxidation with Fer-1 or the mitochondrial ROS – apoptosis cascade with ZVAD had no effect on BBB integrity. Since a ferroptosis-like process was induced, we turned our attention to the effect on cellular iron homeostasis and found that upon insult the endothelial cells underwent a dramatic increase in their LIP. DFO inhibited this surge and repressed both injury induced permeability and TEER damage. Treatment with DFO abrogated the inhibition in expression of the adherens junction and master regulator of BBB integrity, Ve cadherin. Additionally, DFO increased the expression of the hypoxia driven transcription factor HIF2α. The DFO - HIF2α axis had a direct and exclusive effect on BBB integrity since nullifying HIF2α activity with a specific HIF2α inhibitor abrogated DFO rescue of endothelium functionality but not their viability. Even though DFO rescued Ve-cadherin cellular expression post transcriptionally, HIF2α inhibitor selectively nullified the rescue of Ve-cadherin on the cell surface while having no effect on total Ve-cadherin expression. In addition, we discovered a novel nitroxide, JP4-039, which also had the dual synergetic effect of rescuing both endothelium cell viability and functionality.

Most in-vivo experiments examining chemical toxins effects on BBB have been performed on rodents, especially experiments elucidating ROS and oxidative stress effects on the BBB^68–70^. Yet, rodents BBB differ from that of humans by their permeability to P-glycoprotein substrates ^71^ and expression levels of tight junctions, such as claudin-5 ^72^ and receptors, such as Transferrin Receptor1^73,74^. Since our model is based on human cord blood derived hematopoietic stem cells that have been co-cultured with pericytes to differentiate into BBB endothelium, the conundrum of discrepancies between species has been addressed. Additionally, this in-vitro human BBB model correlated well with human pharmacokinetic parameters^34^. We also have the advantage of the use of non-immortalized cells, since immortalized cell lines have accumulated mutations, mycoplasma contamination, loss of BBB-like phenotype and authenticity complications^75,76^, resulting in varying BBB characteristics between intraspecies endothelial cell lines^77^.

As previously reported, both the apoptosis inhibitor ZVAD and the antioxidant TEMPOL abrogated PX-induced endothelial cell viability damage; yet, they had no significant effect on the reduced barrier functionality^36^. It has been widely reported that ROS affects BBB integrity not only in in-vivo ^69^ and in-vitro^78,79^ models, but that there is also evidence of ROS-induced BBB damage under varying pathologies such as stroke, traumatic brain injury (TBI) and Neurodegenerative diseases ^80–84^. Thus, we decided to elucidate whether a more specific localized oxidative stress was responsible for BBB functional damage. Since membrane phospholipids are sensitive to ROS attacks^85^ and it has been previously reported in other BBB models that lipid peroxidation damages BBB integrity^49^, we measured lipid peroxidation and examined its effects on BBB integrity. A rapid increase in lipid peroxidation was evident upon insult accumulating over 24 hours (Fig 1a-b). It has been previously reported that lipid peroxidation induces ferroptosis, a relatively new non-apoptotic, iron-dependent regulated form of cell death caused by the accumulation of lipid-based ROS^52^. Even though there have been several studies in neurons on this process^86–88^, no study to date has been undertaken to elucidate its role in BBB endothelium. To ascertain whether this programmed cell death was activated in our model, we added both the classic ferroptosis and lipid peroxidation inhibitor; Ferrostatin-1 and the iron chelator DFO. Both compounds attenuated cell viability damage (Fig 1c). To validate ferroptosis, the absence of apoptosis needs to be confirmed. Even though caspase-3 activation was recorded in conjunction with significantly increased cell death 10 hours post insult (Fig 2d), Fer-1 had no effect on apoptosis (Fig 1d). These findings negate the possibility of the classic ferroptosis being induced since ultimately the cells underwent apoptosis. Recent reports have highlighted ferroptosis ability to sensitize the cell to compound-induced apoptosis, such as the tumor necrosis factor related apoptosis-inducing ligand (TRAIL). Ferroptosis induces the expression of the apoptosis modulator p53 upregulated modulator of apoptosis (PUMA), yet this modulator alone does not initiate apoptosis, but rather exacerbates an already active process^89^. Therefore, a similar chain of events can explain the role of ferroptosis in our model. We wanted to elucidate the downstream cascade that results in apoptosis and clarify how ferroptosis could play a part in this process. Apoptosis essentially consists of two different pathways; the intrinsic and extrinsic pathways. In the intrinsic, or mitochondrial pathway of apoptosis, effector caspases are activated after a signaling cascade involving mitochondrial outer membrane permeabilization, causing the release of cytochrome c and the activation of caspase-3 that orchestrates the death of the cell^90^. In the extrinsic, or death receptor pathway of apoptosis, ligation of death receptors on the cell surface leads to the formation of a death-inducing signaling complex, which includes FADD and caspase-8^91^. We showed that the classic intrinsic apoptosis pathway exists in our model since insult induced mitochondrial superoxide formation (Fig 2a), mitochondrial membrane permeation (Fig 2b) and caspase-3 activation (Fig 2d). Concordantly, all these processes were inhibited with ZVAD (Fig 2a-d). Although ZVAD abrogates apoptosis, it exacerbates damage to BLEC functionality36 indicating that the intrinsic pathway is not directly responsible for the damage to BBB integrity. The extrinsic pathway appears to also be induced since we have shown in a recent paper that caspase-8 inhibition reduced cell toxicity^37^. As mentioned earlier, unlike Fer-1, the iron chelator DFO rescues BLEC from both cell death and apoptosis (Fig 3a and c). Even though DFO inhibits superoxide formation in the mitochondria (Fig 3f) it seems to inhibit apoptosis through the extrinsic pathways since it exacerbates mitochondrial membrane permeation (Fig 3d), an essential step in the intrinsic pathway.

Interestingly, tumor necrosis factor α (TNF-α) has been reported to induce apoptosis through both intrinsic and extrinsic pathways in head and neck squamous cell carcinoma^92^, and iron accumulation in human umbilical endothelial cells (HUVEC)^93^. Furthermore, DFO inhibits TNF-α induced cytotoxicity in HUVEC^94^, while the iron chelator 1,2-dimethyl-3-hydroxypyridin-4-one inhibited the expression of the TNF-α-induced inflammatory gene; monocyte chemoattractant protein (MCP)-1^95^, a chemoattractant we recently found to be activated in the mouse brain’s vasculature after exposure to PX^37^.

Such evidence indicates the possibility that in our model both apoptosis pathways are induced by TNFα or other yet undefined molecule with similar effects, and that DFO significantly attenuates TNFα signaling or expression, resulting in BLEC viability rescue.

Even though we had elucidated the apoptotic pathways and their kinetics, the process responsible for damage to barrier functionality still eluded us. We turned our attention to possible changes in iron since ferroptosis has been reported to deregulate iron homeostasis through manipulation of iron regulatory proteins such as ferritin and transferrin^96,97^. Additionally, ferroptosis-induced degradation of ferritin resulted in an accumulation of the cellular liable iron pool^96^.

Indeed, a rapid and dramatic increase in the liable iron pool was detected upon insult, while DFO slightly yet significantly reduced this upsurge (Fig 3b). Iron overload is associated with loss of BBB integrity in an in-vivo model of transient forebrain ischemia in rats^98^ and has been reported to induce endothelial cell toxicity in an in-vitro model of secondary brain injury after brain hemorrhage^64^. We found that DFO not only rescued BLEC viability (Fig 3a) but also reduced permeability and restored TEER (Fig 3g). Furthermore, the addition of FAS to induce LIP resulted in damage to BBB integrity, yet did not significantly affect BLEC cell viability (Fig 4d), demonstrating that a deregulation in the endothelial iron homeostasis might be crucial for BBB functionality with no effect on cell viability.

To elucidate the mechanism by which DFO rescues BBB integrity, we analyzed the expression of the adherens junction protein Ve-cadherin. DFO abrogated Ve-cadherin decrease in protein expression (Fig 4a) but not mRNA expression (Fig 4e), indicating a post transcriptional Ve-cadherin rescue. To find a link between DFO and Ve-cadherin, we turned our attention to the iron regulated HIF2α. HIF2α has been reported to regulate Ve-cadherin expression directly by targeting its promoter in cardiac endothelial cells63 or indirectly by targeting the promoter of VE-PTP in lung microvascular endothelial cells. Ve-PTP stabilizes Ve-cadherin by preventing its internalization and degradation^62^. Yet the role of HIF2α in brain endothelial cells and BBB permeability has not been discovered.

We found here that DFO significantly increased HIF2α expression 24 hours post insult (Fig 4b). To establish the regulatory role of HIF2α on BBB integrity we applied a specific HIF2α inhibitor, that exclusively targets HIF2α but not HIF1α^99^. This inhibitor abrogated both DFO permeability and TEER rescue while having no significant effect on either DFO inhibition of cell toxicity (Fig 4c) or mitochondrial superoxide formation (Fig 4h), strengthening our hypothesis that BLEC viability and integrity are independently regulated by two separate pathways. To validate the role of the iron liable pool in damage to BBB functionality through HIF2α expression, we replaced PX with the classic liable iron pool inducer ferrous ammonium sulfate^64^. While endothelium toxicity was not significantly affected, damage to BBB functionality was evident by enhanced permeability and reduced TEER. This damage was reversed by DFO, while HIF2α abrogated the effect of DFO (Fig. 4d). These results reiterated our theory that the barrier integrity is regulated by iron homeostasis in BLEC through the HIF2α – Ve-cadherin axis (Fig. 6). Surprisingly, HIF2α inhibitor had no significant effect on Ve-cadherin mRNA or total cell protein expression (Fig 4e and f) but exclusively abrogated DFO rescue of Ve-cadherin expression on the cell surface (Fig 4g). A similar phenomenon has been recently reported with TNFα and its effect on Ve-cadherin cell surface expression in HUVEC^100^. This supports our hypothesis that TNFα is involved in the mechanisms of our model. Further studies should be made to elucidate the role of TNF-α in both brain endothelium viability and integrity in our injury model.

**Fig. 6.**
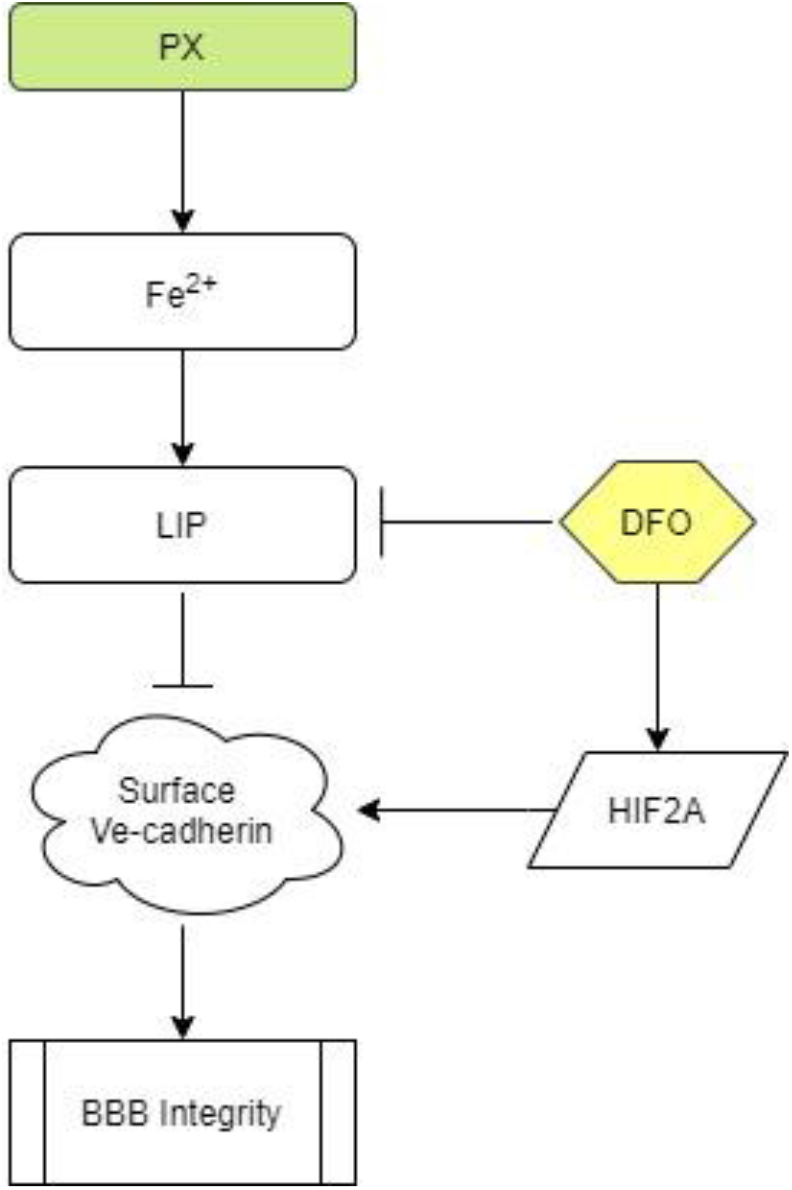
BBB integrity is regulated by the iron dependent HIF2α-Ve cadherin axis. Flowchart elucidating the molecular pathways that regulate BBB integrity in our model. Upon insult the liable iron pool is significantly increased, resulting in damage to the BBB integrity. DFO rescues BBB functionality by inducing the expression of HIF2α that results in the restoration of Ve-cadherin on the cell surface.

It has long been established that acute hypoxia induces BBB permeability^101,102^. Conversely, moderate chronic hypoxia has been shown to enhance BBB stability through the expressions of laminins^103^. HIF1α as opposed to HIF2α has a negative correlation to BBB integrity^104,105^. This can be explained by their opposing roles; HIF1α governs the acute adaptation to hypoxia, whereas HIF2α activity is induced later^106^, and this creates a transitional switch between the expression of the HIF1α acute hypoxia targeted genes to the expression of the HIF2α chronic hypoxia targeted genes. The present discovery of a novel HIF2α – Ve-cadherin brain endothelium axis that regulates BBB integrity through iron homeostasis can shed light on the molecular mechanisms involved in the maintenance of BBB integrity upon hypoxia-independent injuries and might furthermore indicate the pathways that are responsible for enhanced BBB integrity in moderate chronic hypoxia. We speculate that laminin could be the missing link in our axis that enables HIF2α to regulate Ve-cadherin surface expression since it has been reported that chronic mild hypoxia increases expression of laminins 111 and 411 and the laminin receptor α6β1 integrin at the BBB^103^. Furthermore, laminin 511 was shown to induce Ve-cadherin localization to cell-cell borders via β1 and β3 integrins^107^.

Further studies are needed to elucidate the role of laminins and their receptors and to gain a deeper understanding of the molecular mechanisms of HIF2α regulated BBB integrity in our model.

Iron homeostasis balance is an essential component of cellular physiology and is a delicately balanced process. On the one hand, it is involved in critical intracellular processes, including DNA synthesis and cellular respiration. In contrast, iron overload results in ROS formation and oxidative stress through the Fenton reaction ^108–110^. We therefore sought to find another compound, which will not act as a classic chelator. We have synthesized and screened more than 30 new compounds (not shown) and identified a nitroxide, JP4-039–which is a mitochondrial targeted antioxidant and free radical scavenger^111,112^ – as a potential candidate. It has been demonstrated that JP4-039 in combination with a bone marrow transplant, increased tight junction protein expression and barrier integrity in epithelial intestinal cells of mice who had been exposed to total body irradiation without affecting cell viability^113^. JP4-039 had multifaceted beneficial effects in our BBB injury model; it inhibited mitochondrial superoxide formation, rescued BLEC cell death via apoptosis inhibition and most importantly, it protected BBB integrity by attenuating BBB increased permeability and TEER decline (Fig 5). The precise molecular mechanism of JP4-039 still needs to be elucidated; nevertheless, we have identified a novel non-toxic chemical that both protects BLEC viability and integrity, with the potential of incorporating it as a therapeutic compound in treatments of diseases associated with BBB breakdown, such as Neurodegenerative diseases.

In conclusion, using an established human BBB model, we have shown that chemical insult rapidly induces lipid peroxidation and a ferroptosis-like event. This event potentially enhances separate pathways that result in apoptosis. We have elucidated that both intrinsic and extrinsic apoptosis pathways are present in our model and we have shown the kinetics of these pathways. More importantly, we have demonstrated that the source of BBB breakdown in our model is not a direct result of ROS related processes but rather an imbalance induced in the cellular iron homeostasis of BBB endothelium. We have furthermore elucidated regulation of BBB integrity through the iron dependent HIF2α – Ve-cadherin pathway and that the FDA approved DFO used to treat acute iron poisoning and hemochromatosis in clinic^114^, attenuates damage to BLEC viability and barrier integrity by upregulating this pathway. Since there is a high correlation between cognitive decline, Neurodegenerative diseases and exposure to organophosphate^39,115–117^, and several in vivo studies have shown that BBB integrity is compromised after exposure to organophosphates^118^ and specifically to PX^42^, we have used PX to induce BBB damage in an attempt to shed light on the mechanisms of BBB breakdown which might be generalized– with proper cautious– to other brain pathologies in which the BBB is compromised. These results can help clarify the involvement of iron in BBB breakdown and Neurodegenerative diseases since it has been reported that BBB dysfunction is an early biomarker of Neurodegenerative diseases7 and that there is a clear correlation between iron overload and Neurodegenerative diseases^119^. Additionally, it can help solve the conundrum of iron’s involvement in initiating the events that result in Neurodegenerative diseases. Moreover, a novel compound, JP4-039, was found to treat BBB breakdown. These findings are of critical importance since BBB breakdown is a predominant factor in many neurological diseases. Our research has elucidated the mechanisms that drive both endothelium viability and functionality under PX injury. We expect that these findings can be used to assist in developing a viable treatment to attenuate BBB damage, thereby inhibiting BBB-associated disease progression.

## Materials and Methods

### Reagents

Mouse anti-VE-cadherin antibody (sc-6458) diluted 1:50 was obtained from Santa Cruz Biotechnology (United States). Cy and Alexa Fluor-conjugated secondary antibodies were acquired from Jackson Immunoresearch (United States) and Molecular Probes (United States), respectively, and used for immunocytochemistry. Z-VAD-FMK was obtained from Adooq-bioscience (187389-52-2, United States) and Santa Cruz Biotechnology (sc-3067, United States). Marimastat from Santa Cruz Biotechnology (sc-202223, United States), MG-132 from Promega (G9951, United States). Cytotoxgreen was obtained from Essen BioScience (United States) and CellROXgreen from Molecular Probes (United States). PX-ethyl was purchased from Sigma (United States), according to Sigma safety data sheet safety measures of eyeshields, face shields, full-face respirator and Gloves should be taken. For assessing the BBB response, PX freshly made in ethanol to 400 mM stock solution was immediately diluted in the medium to the desired final concentrations and added to the cell culture. All other reagents applied in this study were used in accordance to the known literature and the supplier’s guidelines.

### Media

Brain-like endothelial cell and pericytes were grown in ECM medium (Sciencell, United States) that was composed as follows: 5% fetal calf serum (Gibco, United States), ECGS supplements and 50 mg/ml gentamicin (Biological industries, Israel).

### BBB in-vitro model

To investigate the cellular response to chemical injury, a human BBB model which is well characterized for studying BBB injury, inflammation and barrier functionality was utilized^34,36^. The generation of these ECs relies on biological principles observed in the repair of BBB in the human body. The in-vivo repair of the endothelium is mediated by endothelial progenitor cells that migrate to the sites of endothelial injury, incorporate in the growing vessel and differentiate into ECs^120,121^. The human BBB model was generated using human cord blood-derived hematopoietic stem cells; CD34+ cells were isolated from umbilical cord blood. Parents’ infants signed a consent form. All protocols were done with the authorization of the French Ministry of Higher Education and Research (CODECOH Number DC2011-1321). CD34+ cells were initially differentiated into ECs followed by the induction of BBB properties by co-culture with bovine brain pericytes on matrigel-coated insert (Transwell^®^, 5μm). The produced BLECs express TJ proteins and transporters typically observed in brain endothelium and maintain expression of most in-vivo BBB properties for at least 20 days^34^. We used the co-culture system or monoculture of CD34+ ECs grown in PCM. BBB injury was generated by exposure to the organophosphate PX, as previously characterized^36^.

### Cells

Human CD34C-derived ECs and bovine brain pericytes were obtained from Artois BBB laboratory where their isolation and differentiation were conducted as previously described^34,122,123^. Regarding the collection of human umbilical cord blood: infants’ parents signed an informed consent form, in compliance with the French legislation. The protocol was approved by the French Ministry of Higher Education and Research (CODE-COH Number DC2011-1321). All experiments were carried out in accordance with the approved protocol. For each experiment, the cells were expanded on gelatin (Sigma, United States)-coated dishes in ECM medium. For BLEC monolayer experiments, CD34C ECs were cultured with pericyte-conditioned-medium for 4 days. For co-culture experiments, 5×10^4^ brain pericytes were seeded on 12 well gelatin-coated plates (Costar, Corning, United States) and cultured in ECM medium. Human CD34C-derived ECs were seeded at a density of 8×10^4^/insert onto the Matrigel-coated (BD Biosciences, United States) Transwell inserts (3401-Costar, Corning, United States). Cells were grown in co-culture for 6–8 days for Pe assay and acquire BBB properties during this period becoming BLECs (BLECs).

### Permeability Assay

Prior to the experiments, HEPES-buffered Ringer’s solution (RHB) was added to empty wells of a 12 well plate (Costar). Filter inserts, containing confluent monolayers of BLECs on the luminal side and pericytes on the abluminal side were placed in the 12 well plate filled with RHB to acclimate for 30 min at 37°C, after which compound solution containing the fluorescent integrity marker Fluorescein (50 mg/ml; Sigma, United States) was added to the luminal side, and then placed on a shaker at 37°C. Every 10 min inserts were transferred to a new 12 well plate over a period of 40 min. Aliquots from the abluminal solution were taken from each time point and the fluorescence was quantified. Inserts without cells were tested in each Pe measurement. Fluorescein detection was carried out on an Infinite 200 PRO (Tecan, Switzerland) plate reader using the excitation/emission wavelength (nm) settings: 485/538. Pe coefficient was obtained from the slope of the calculated clearance curve as described in reference^124^. Typical Pe value for the control was Pe = 0.35×10 ^−3^ cm/min.

### TEER Assays

Impedance spectrum measurements were performed using a multi-well impedance spectrometer (cellZscope, Nano analytical, Germany). Filter inserts, containing confluent monolayers of BLECs on the luminal side and pericytes on the abluminal side were placed in the cellZscope and impedance was measured at an hour interval for 12 h prior to TW treatment and 24 h post treatment while incubated at 37°C and 5% CO2.

The modulus (|Z|) and the phase (θ) of the impedance were obtained for each treatment. TEER is then extracted from the model and multiplied by the membrane area (1.12 cm^2^) and is presented in Ώ.cm^2^. TEER values in the current study reached an average of 30 ± 5 Ώ.cm^2^ before treatments were applied.

### Cell Death by LDH Release

The toxicity of PX was investigated on monolayers of BLEC seeded on 96 well plates or on inserts luminal side in co-culture experiments using the commercially available Cytotoxicity Detection Kit (Promega, United States). An aliquot of 25 μl medium was taken to quantify the lactate dehydrogenase (LDH) release. The test was performed according to the manufacturer’s instructions and absorption was measured at 490 nm by an Elisa plate reader (Tecan, Switzerland).

### Cell Death by Cytotoxgreen Staining

Monolayers of BLEC were treated with PX in 96 well plates and monitored simultaneously for cytotoxic response kinetics with cytotox green stain (250 nM) for 24 h by live imaging using an IncuCyte imaging system (Essen BioScience, United States). Images were captured every hour and cells with a positive fluorescence signal were counted by the IncuCyte^®^ integrated analysis software.

### Oxidative Stress Analysis

Cellular oxidative stress was detected with the use of the cell-permeable fluorogenic probe CellROX (5 mM; Molecular Probes, United States). Monolayers of BLECs were treated in 96 well plates and monitored with CellROX stain for 24 h by live imaging using an IncuCyte imaging system (Essen BioScience, United States). The intensity of CellROX fluorescence was calculated and analysed by the IncuCyte^®^ integrated analysis software to quantify ROS levels.

### Apoptosis Analysis

Apoptosis was measured with the use of the Apoptosis cell-permeable non-toxic fluorogenic Caspase-3/7 Reagent (Essen BioScience, United States). Monolayers of BLECs were treated in 96 well plates and monitored with Caspase-3/7 Reagent for 24 h by live imaging using an IncuCyte imaging system (Essen BioScience, United States). Images were captured every hour and cells with a positive fluorescence signal were counted by the IncuCyte^®^ integrated analysis software.

### Mitochondrial ROS Formation Analysis

Superoxide formation in the mitochondria was measured with the use of the MitoSOX™ Red reagent (Molecular Probes, United States). MitoSOX™ is fluorogenic dye specifically targeted to mitochondria in live cells, oxidation by superoxide exclusively produces red fluorescence. Monolayers of BLECs were treated in 96 well plates and monitored with MitoSOX™ for 24 h by live imaging using an IncuCyte imaging system (Essen BioScience, United States). Images were captured every hour and the fluorescence signal intensity was calculated and analysed by the IncuCyte^®^ integrated analysis software to quantify mitochondrial ROS formation levels.

### Mitochondrial Membrane Depolarization Analysis

Mitochondrial Membrane Depolarization was measured with the use of the membrane-permeant JC-1 dye (Molecular Probes, United States). JC-1 aggregates in the mitochondria emitting a red fluorescence signal (emission of 590±17.5 nm) and exists as a monomer in the cytoplasm emitting a green fluorescence signal (emission of 530±15 nm) as a result of mitochondrial membrane depolarization.

Monolayers of BLEC were treated in 96 well plates and monitored with the JC-1 dye for 24 h by live imaging using an IncuCyte imaging system (Essen BioScience, United States). Images were captured every hour and the increase in the green-red intensity fluorescence signal ratio was calculated and analysed by the IncuCyte^®^ integrated analysis software to quantify mitochondrial membrane depolarization.

### Lipid peroxidation Analysis

Lipid peroxidation was measured with the use of C11-BODIPY^581/591^ (Molecular Probes, United States). This probe incorporates readily into cellular membranes and upon oxidation, the fluorescence of this fluorophore shifts from red (~590 nm) to green (~510 nm). The fraction of oxidized C11-BODIPY581/591 is calculated by dividing the green fluorescence intensity by the red fluorescence intensity signal based upon the protocol previously reported^125^. Monolayers of BLECs were treated in 96 well plates and monitored with C11-BODIPY^581/591^ for 24 h by live imaging using an IncuCyte imaging system (Essen BioScience, United States). Images were captured every hour and the increase in the green-red intensity fluorescence signal ratio was calculated and analysed by the IncuCyte^®^ integrated analysis software to quantify lipid peroxidation. Additionally, the lipid peroxidation assay kit (Abcam, United States) was also used according to the manufacturer protocol.

### Calcein-AM analysis

Cellular liable iron pool (LIP) levels was measured with the use of calcein acetoxymethyl ester (CA-AM) (Sigma-Aldrich, St. Louis, MO). CA-AM enters viable cells and becomes fluorescent upon hydrolysis by esterases; its fluorescence is quenched by binding of LIP^126–128^. Monolayers of BLECs were treated in 96 well plates and simultaneously incubated with CA-AM (1 *μ*M) (Sigma, St. Louis, MO). Following staining, cells were analyzed by live imaging using an IncuCyte imaging system (Essen BioScience, United States). Images were captured every hour and the increase in the green intensity fluorescence signal was calculated and analysed by the IncuCyte^®^ integrated analysis software to quantify relative LIP concentration.

### Western blot Assay

BLEC were grown on gelatin-coated 6-well plates and treated for 24 hours. The cells were then lysed with RIPA Buffer (Sigma, United States) and protease inhibitors (Roche, France). Cell extracts were separated by 10% SDS–PAGE followed by blotting on nitrocellulose (Whatmann Schleicher-Schuell, France) and analyzed with a mouse anti-Ve-cadherin (Santa Cruz Biotechnology, United States), and normalized with a rabbit anti-actin (Santa Cruz Biotechnology, United States) antibodies.

### Immunocytochemistry

CD34C-derived BLEC were grown on gelatin 96-wells and treated for 24 hours. After treatments, the cells were fixed with 4% paraformaldehyde (PFA) for 10 min at room temperature and exposed to blocking solution [10% horse serum/ 0.1% triton/ (PBS)] for 1.5 h. The BLEC were then incubated with anti-VE-cadherin and anti-HIF2α (overnight, 4C֯), washed with PBS/0.1% Tween20 and immunostained with appropriate secondary antibodies (1 h, room temperature). Cells were counterstained with propidium iodide (PI) (5 min). Images were taken with IncuCyte^®^ fluorescence microscope (Essen BioScience, United States) with 10objective. Antibody intensity fluorescence signal was calculated and analyzed by the IncuCyte^®^ integrated analysis software and normalized by cell count using the PI signal. When comparison between different treatments was made, the exact same optical settings were used to avoid any misinterpretation of the results. The brightness of some pictures in this paper was increased artificially to emphasize the pattern of protein expression.

### Gene expression – rtPCR

BLEC were grown on gelatin-coated 6-well plates and treated for 24 hours. RNA isolation was performed using the RNA Purification kit (NucleoSpin^®^) following the protocol of purification from cultured cells and tissues. The cDNA was synthesized using the qScript™ cDNA Synthesis Kit (Quanta Bioscience™). To precisely quantify specific mRNA expression, RT-PCR Step One Plus system (#8024, Applied Biosystems) was used.

The following PCR-program was performed: 20s 95°C (initial denaturation); 95°C for 15s, 60°C for 30s, repeated 40 times (amplification); 95°C for 15s, 60°C for 1min, 95°C for 15s (melting curve). The PCR results were evaluated using the Applied Biosystems software.

Primers used:

Ve-cadherin:

Fw-5′-GGTCCCTGAACGCCCTGGTAA-3′
Rev-5′-GGAGTGGAGTATGGAGTTGGAGCA-3′

GAPDH as housekeeping gene:

Fw-5′-GGCCTCCAAGGAGTAAGACC-3′
Rev-5′-AGGGGTCTACATGGCAACTG-3′

### Flow cytometry (FACS)

BLEC were grown on gelatin-coated 6-well plates and treated for 24 hours. After treatment BLEC were collected using Cell Dissociation Non-enzymatic Solution (Sigma, United States), washed, and suspended in cold PBS. One million (1 × 10^6^) cells were incubated with Anti-Ve cadherin Alexa Fluor 488 conjugated or isotype control (mouse IgM) (Invitrogen, United States) for 40 min on ice. After three washes with cold PBS, cells were examined in the CyFlow^®^ Cube 6 (Sysmex America, United States) and data was analyzed using the FlowJo™ v10.7 software (BD biosciences, United States).

### Data and statistical analysis

Results are expressed as the mean ± standard deviation of the mean, with at least three biological repeats unless mentioned otherwise. Comparison between three groups or more was analyzed using One-way analysis of variance (ANOVA) with Tukey's multiple comparison test for post-hoc analyses. Live cell imaging data was analyzed with Two-way ANOVA with Tukey's multiple comparison test for post-hoc analyses. GraphPad Prism 8.0 software was used for all statistical analysis (GraphPad Software Inc., La Jolla, CA). Differences were considered significant at P values <0.05.

